# Label-free citrate-stabilized silver nanoparticles-based highly sensitive, cost-effective and rapid visual method for differential detection of *Mycobacterium tuberculosis* and *Mycobacterium bovis*

**DOI:** 10.1101/2023.08.04.551932

**Authors:** Naresh Patnaik, Ruchi Jain Dey

## Abstract

Tuberculosis poses a global health challenge, demanding improved diagnostics and therapies. Distinguishing between *Mycobacterium tuberculosis* (*M. tb*) and *Mycobacterium bovis* (*M. bovis*) infections holds critical “One Health” significance due to zoonotic nature of these infections and inherent resistance of *M. bovis* to pyrazinamide, a key part of Directly Observed Treatment, Short-course (DOTS) regimen. Furthermore, most of the currently used molecular detection methods fail to distinguish between the two species. To address this, our study presents an innovative molecular-biosensing strategy. We developed a label-free citrate-stabilized silver nanoparticle aggregation assay, offering sensitive, cost-effective, and swift detection. For molecular detection, genomic markers unique to *M. tb* and *M. bovis* were targeted using species-specific primers. In addition to amplifying species-specific regions, these primers also aid detection of characteristic deletions in each of the mycobacterial species. Post polymerase chain reaction (PCR), we compared two highly sensitive visual detection methods with respect to the traditional agarose gel electrophoresis. The paramagnetic bead-based bridging flocculation assay, successfully discriminates *M. tb* from *M. bovis* with a sensitivity of ~40 bacilli. The second strategy exploits citrate-stabilized silver nanoparticle which aggregates in the absence of amplified dsDNA on addition of sodium chloride (NaCl). This technique enables precise, sensitive and differential detection of as few as ~4 bacilli. Our study hence advances tuberculosis detection, overcoming challenges of *M. tb* and *M. bovis* differentiation offering a quicker alternative to time-consuming methods.

## Introduction

Tuberculosis (TB) is a chronic infectious disease caused by *Mycobacterium tuberculosis complex* (MTBC) species. In humans, *M. tb* is the major causative agent, while in cattle, *M. bovis* cause the majority of the bovine TB (bTB). A recent global meta-analysis suggests ~12% prevalence of *M. bovis* of the total MTBC cases in human. *M. bovis* is more prevalent in case of extra-pulmonary TB (EPTB) infections compared to pulmonary TB (PTB) ^1–4^. *M. tb* and *M. bovis* pose a significant zoonotic risk. Recent studies provide robust evidence for cattle-to-human and human-to-cattle transmission of MTBC organisms due to close interaction between them, especially in the rural, agricultural and farm settings ^1–4^. Hence, a multisectoral One Health approach is needed for timely detection of *M. tb* and *M. bovis* in point-of-care (POC) settings for effective TB control in human and cattle.

In absence of an effective vaccine and prevalence of multi-drug resistant strains of MTBC, TB continues to ravage humanity causing more than 10 million new infections and ~1.5 million deaths each year (Global Tuberculosis Report, 2022, WHO). Almost, ~66% of the total active TB cases are concentrated in just eight countries (India, Indonesia, China, Philippines, Pakistan, Nigeria, Bangladesh, and South Africa) which have agrarian economy with majority of the population residing in resource poor rural areas (Global Tuberculosis Report, 2022, WHO). Such large number of TB cases put immense burden on the already overwhelmed healthcare system of these nations, further jeopardizing the TB control. Effective management of TB depends heavily on prompt and early diagnosis of TB, recognition of drug resistance profile, robust contact tracing, ensuring patients’ adherence and response to 6-9 months of drug regimen, and continuous screening of TB infection in high-risk groups (Global Tuberculosis Report, 2022, WHO). However, successful achievement of the above listed goals needs cost-effective, fast, simple, sensitive and specific detection tools that can be easily implemented in urban and rural, resource-poor settings in a POC manner.

Among the inexpensive and fast methods of detection, sputum smear microscopy is one of the most common methods utilized in resource-poor settings to detect MTBC which are acid-fast bacilli (AFB). However, the detection sensitivity is ~5,000 to 10,000 bacilli/millilitre (ml) of sputum. Hence, microscopy-based detection may lead to false negative cases causing increase in transmission due to lack of timely detection. A recent study compared sensitivity and specificity of *M. tb* detection in EPTB employing a variety of currently used clinical diagnostic methods and observed only a 25% sensitivity in case of AFB microscopy compared to the state-of-the-art methods, such as culture-based detection (72.7%), Xpert MTB/RIF (85.6%), and MTBDRplus assay (79.4%) with a 100% specificity for all the methods ^5^. Culture test, the current gold standard is highly sensitive and specific and can detect as few as 10-100 viable bacilli/ml of the specimen and can detect both PTB, EPTB, and paucibacillary (low bacterial load) cases, which may not be detected by sputum smear microscopy. This method is also ineffective for diagnosing latent TB infections ^6^. However, culture test allows detection of drug resistance profile, thereby aiding decision to decide the combination of anti-TB drugs to be used. These culture-based tests though, are extremely time-consuming (4-6 weeks), causing massive delays in starting the treatment leading to increased risk of disease transmission. Moreover, it requires specialized biosafety laboratory infrastructure and highly skilled personnel, increasing the overall cost of diagnosis, further complicating diagnosis in resource-poor settings. To address these disadvantages, molecular diagnostic tools are being adopted which are much faster, and more sensitive, though culture-based methods remain highly relevant for drug susceptibility testing and epidemiological studies.

Molecular detection techniques like PCR has resulted in significant advancements in the detection of MTBC ^4^ offering significant advantages over microscopy and culture-based methods for TB diagnosis, epidemiology, and strain identification ^7, 8^. Traditional PCR and recently developed digital droplet PCR are much faster and more sensitive, capable of detecting even low levels of mycobacterial DNA, enabling early and accurate detection of both active and latent TB infections or asymptomatic individuals with clear chest X-ray and negative AFB ^9^. PCR allows rapid screening of large populations, aiding identification and control of outbreaks more efficiently than the traditional methods and is found to perform better than Xpert MTB/RIF in case of paucibacillary TB ^10–12^. Agarose gel electrophoresis is typically used to detect the PCR products in a cost-effective manner compared to more advanced and quantitative methods like real-time quantitative PCR (qPCR), however the former suffers from poor sensitivity ^13, 14^.

The differential detection of *M. tb* and *M. bovis* in clinical settings is critical because the latter is inherently resistant to pyrazinamide (PZA) due to natural mutation in *pncA* gene encoding pyrazinamidase/nicotinamidase ^15^. As PZA is an important part of the classical DOTS therapy for TB, mono- or co-infections with *M. bovis* if go undiagnosed, may compromise the complete treatment of these patients. Since, *M. tb* and *M. bovis* are clinically, radiologically, and microscopically indistinguishable and classical biochemical tests to identify *M. bovis* are not regularly performed due to increased usage of automated liquid culture systems, diagnosing *M. bovis* has become extremely challenging. Further, due to high degree of genetic similarity, finding unique markers for differential diagnosis of *M. tb* and *M. bovis* is also challenging. Several molecular identification methods based on genomic sequences corresponding to IS6110, 16S rDNA, 23S rDNA or internal transcribed spacers (ITS), are incapable of distinguishing between the two mycobacterial species ^16–18^. It is only following genome sequencing of different species of MTBC, that evolutionary relationship and genomic regions unique to each of these species were identified ^18, 19^. As genome sequencing in clinical settings is not feasible considering the high cost, a PCR-based differential detection of these MTBC species is essential. In addition, combining PCR with other biosensing techniques such as nanoparticle-based visual DNA-detection may further enhance the accuracy and sensitivity of molecular detection.

In the past few years, several visual (naked eye) detection methods to detect MTBC have been researched such as, gold nanoparticles ^20–23^, magnetic bead-based flocculation ^24–27^, and lateral flow assays ^28–31^. These methods provide rapid, user friendly, and easy-to-interpret results without the need for specialized equipment or training, making them suitable for resource-poor or POC settings or during outbreaks, where timely diagnosis is critical. These methods can be tailored to work with various clinical specimen types, enhancing their versatility in detecting pathogens from different sources such as sputum, blood, urine, or cerebrospinal fluid.

In this study, we developed a novel citrate-stabilized silver nanoparticles-based method for label-free detection of mycobacterial DNA post amplification. We also compare the sensitivity of this method with paramagnetic bead-based DNA purification-cum-visual detection and traditional agarose gel electrophoresis. Additionally, our methods allow highly sensitive and differential detection of *M. tb* from *M. bovis*.

## Results and Discussion

### PCR based molecular identification of *M. tb* and *M. bovis*

For differential detection of *M. tb* and *M. bovis*, we first identified genomic loci unique to each of these mycobacterial species from literature **(Figure 1)**. *M. tb* genome consists of 12.7 kb region which is absent from *M. bovis,* this locus consists of 11 genes spanning from *Rv1966* to *Rv1976* and a small region belonging to *Rv1977* **(Figure 1)** ^32, 33^. We designed primers specific to *Rv1966* (*mce3A*), the first gene in this locus. For *M. bovis,* we designed primers corresponding to *Mb1582* (*mmpS6*) gene present in the 0.537 kb locus called Tb deletion region 1 (TbD1), this locus is unique to *M. bovis* and absent in *M. tb* **(Figure 1)** ^34, 35^. We also designed primers reported for *esxA* gene present both in *M. tb* (*Rv3875*) as well as *M. bovis* (*Mb3905)* ^36–38^. This gene is typically used in molecular detection of MTBC, hence it is used as a positive control in this study. (For primer details see **Table 1**)

**Figure 1.**
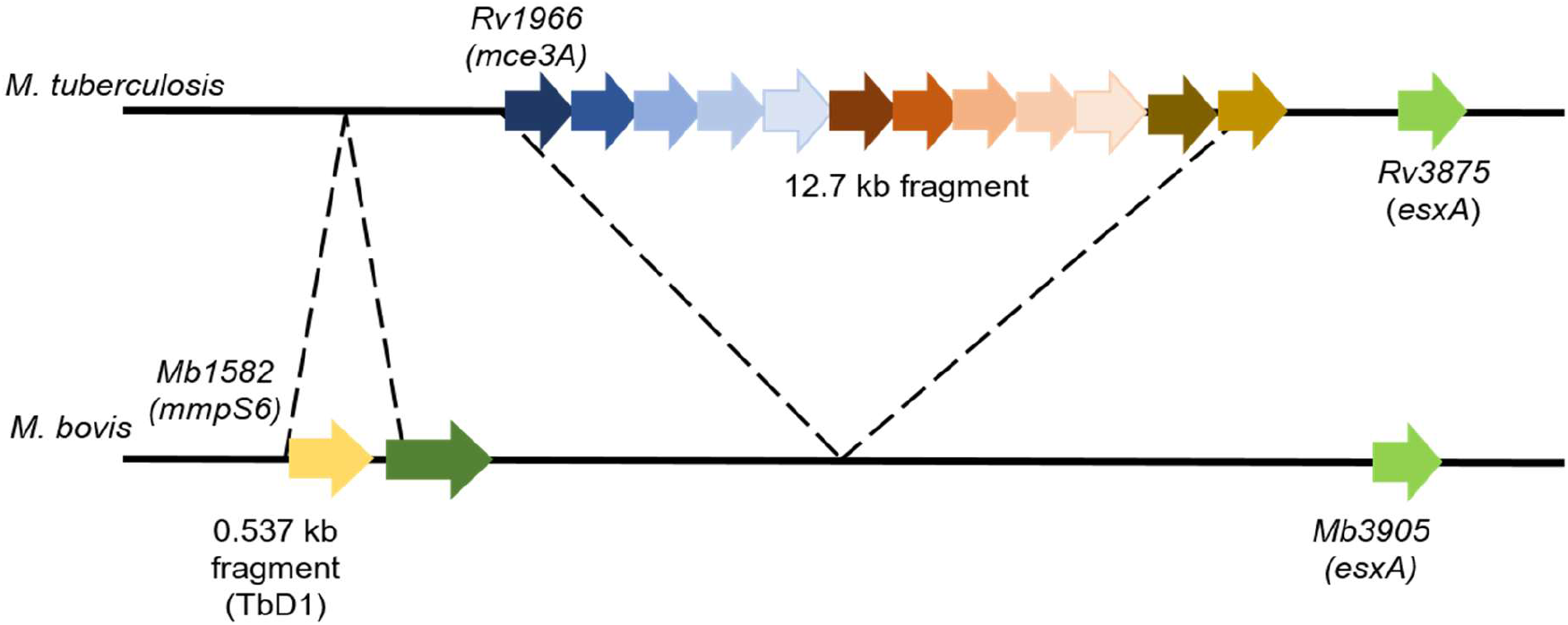
Loci diagram of the target genes used for molecular detection of *M. tuberculosis* and *M. bovis*. A 12.7 kb fragment present in *M. tuberculosis* and absent from *M. bovis* is identified for designing primers corresponding to *Rv1966 (mce3A)* unique to *M. tuberculosis.* Primers corresponding to *esxA* (*Rv3875* in *M. tuberculosis* and *Mb3905* in *M. bovis)* were used as standard method for molecular detection of MTBC. A 0.537 kb region present in *M. bovis* and absent in *M. tuberculosis* called as Tb deletion region 1 (TbD1) was identified for designing primers corresponding *to mmpS6* (*Mb1582*) unique to *M. bovis*.

**Table 1.**
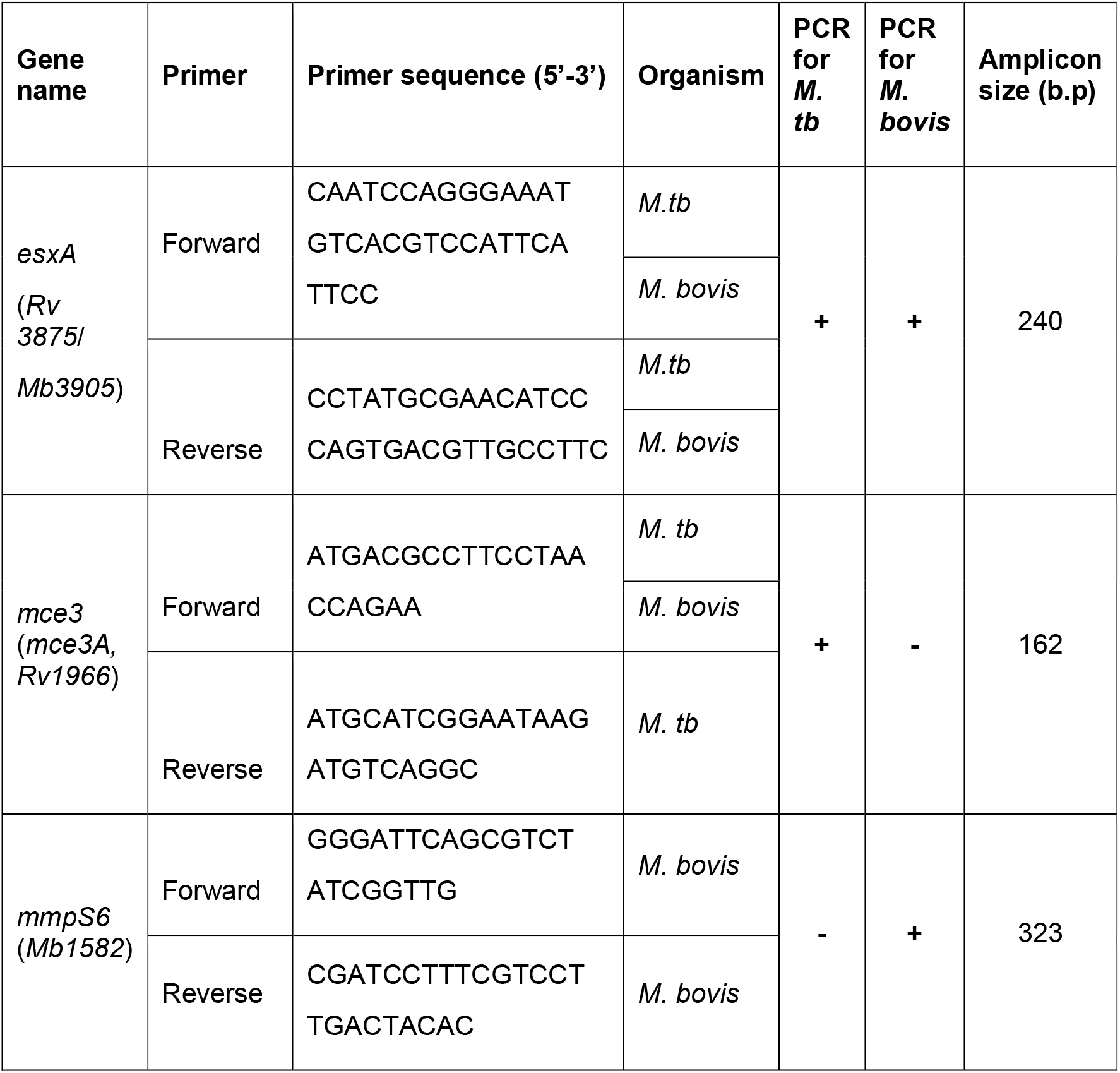
Oligonucleotide primers for the molecular identification of *M. tuberculosis* and *M. bovis*.

The specificity (or cross-reactivity) of the primers designed for *M. tb* and *M. bovis* were determined by carrying out PCR using genomic DNA derived from various organisms including human and important human pathogens such as, *Neisseria gonorrhoeae*, *Chlamydia trachomatis*, *Ureaplasma urealyticum* along with the test organisms, *M. tb* and *M. bovis.* The primers are found to be specific against the test organisms with no cross-reactivity against any of the above listed organisms **(Figure 2 A, B, and C)**.

**Figure 2.**
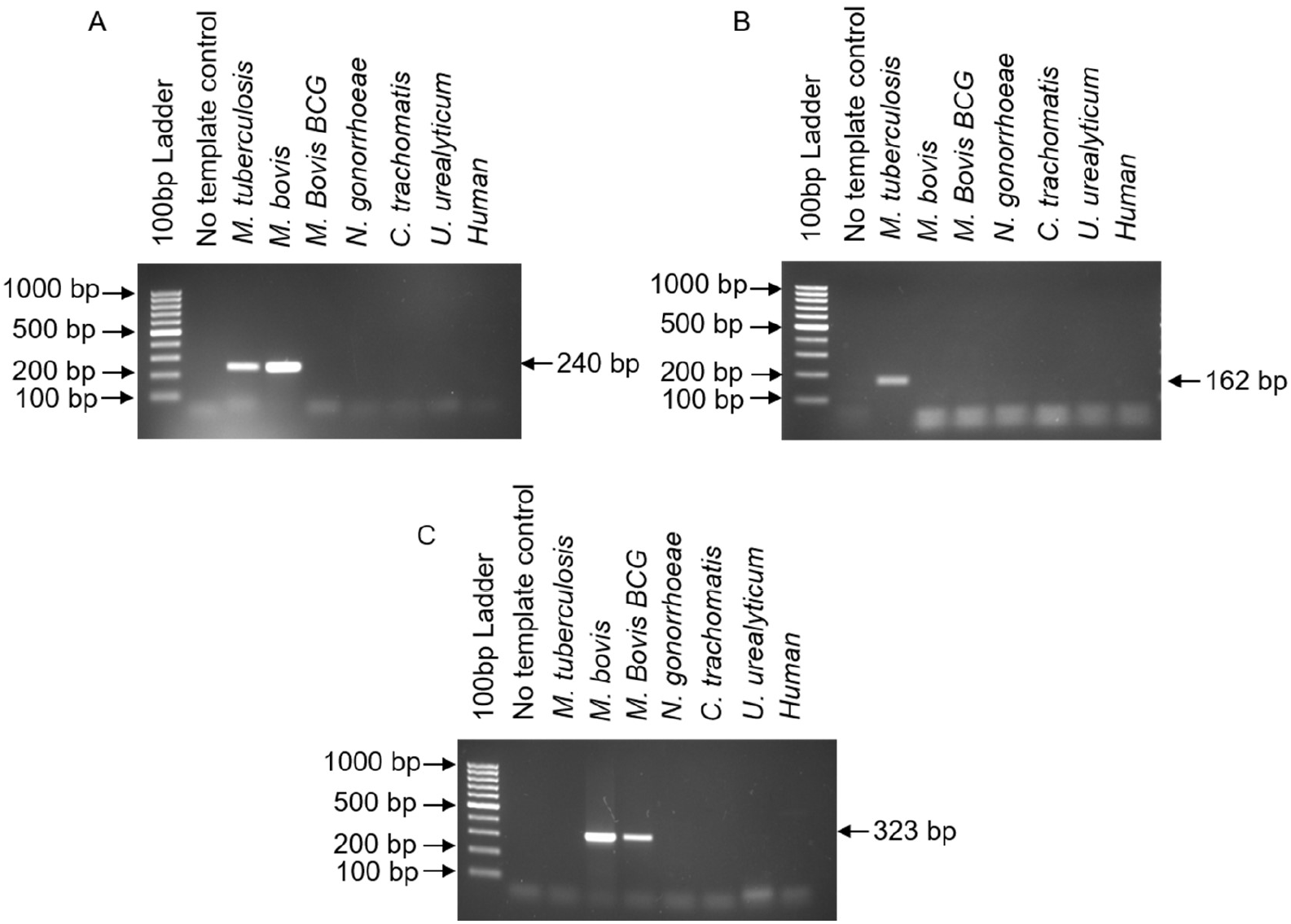
Primer specificity and cross-reactivity verification. Figure depicts agarose gel-based verification of PCR carried out using A. *esxA* gene specific primers with different genomic DNA samples derived from *M. tuberculosis, M. bovis, M. bovis BCG, N. gonorrhoeae, C. trachomatis, U. urealyticum and human. esxA* primers result in a PCR amplicon of 240 bp specific to *M. tuberculosis and M. bovis.* B. *mce3A* gene specific primers result in a PCR amplicon of 162 bp specific to *M. tuberculosis* only. C*. mmpS6* gene specific primers result in a PCR amplicon of 323 bp specific to *M. bovis* and *M. bovis* BCG strain.

### Paramagnetic bead-based bridging flocculation assay for the visual detection of *M. tb* and *M. bovis* in comparison to the agarose gel electrophoresis

In this study, we tested two methods for visual detection of PCR products obtained on gene amplification and compared the sensitivity of detection with respect to the traditional agarose gel electrophoresis. **Figure 3** illustrates how paramagnetic beads can be repurposed for visual detection of the amplified DNA. In addition, the purified products can also be sensed by other cost-effective and simpler visual detection such as silver nanoparticle post-purification with paramagnetic bead as shown later in this study.

**Figure 3.**
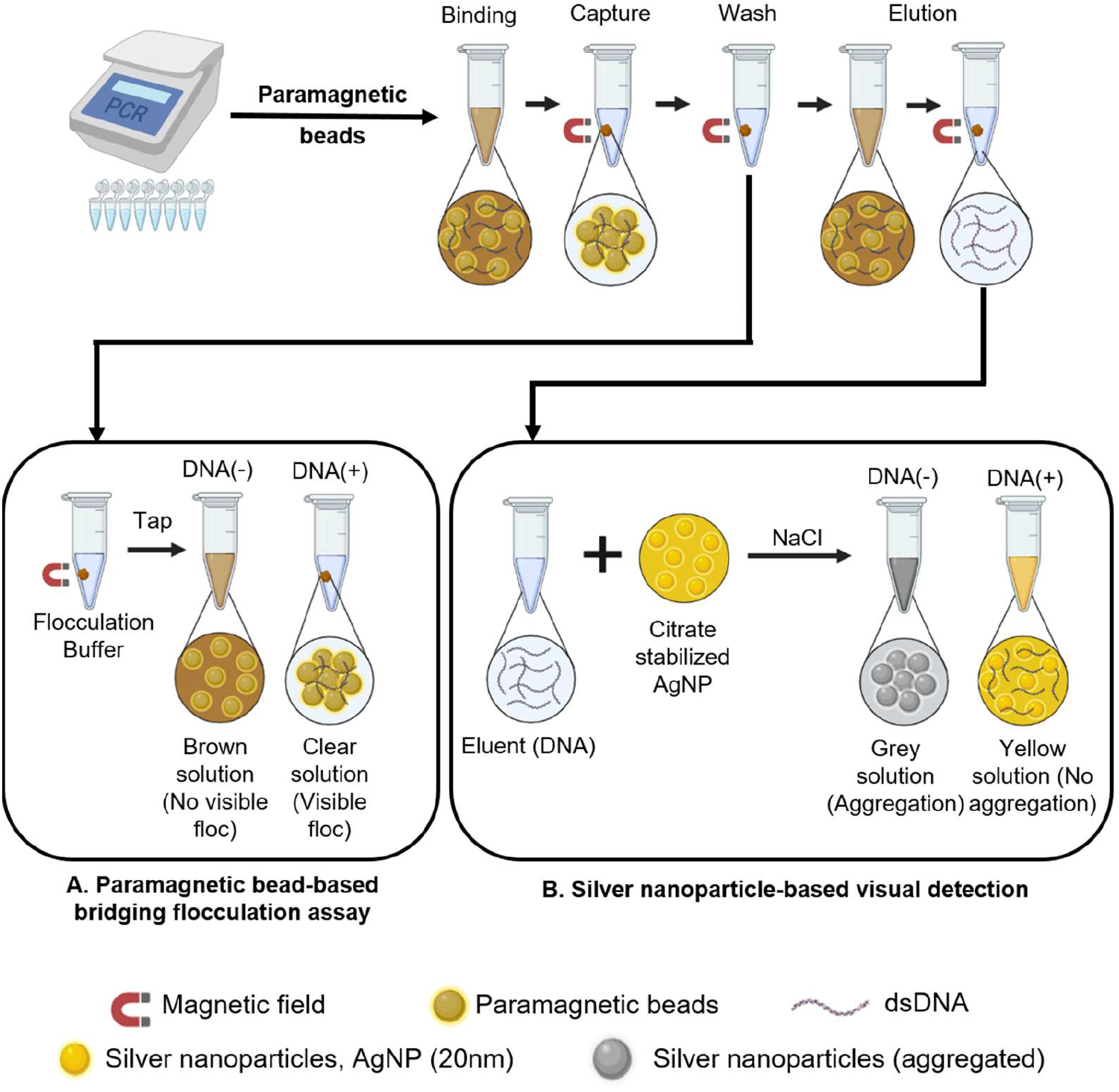
Schematic diagram showing two different types of visual detection. These detection techniques are comparatively more sensitive than the traditional agarose gel-based DNA detection. The amplified PCR products are purified using paramagnetic beads following the steps, namely, binding, magnetic capture, washing and elution. A. The paramagnetic beads can be repurposed for visual detection after the washing step, by using a flocculation buffer, which flocs out DNA adsorbed paramagnetic particles. The paramagnetic particles do not make a floc if sufficient DNA is not adsorbed onto their surface. B. After the elution step, citrate-stabilized silver nanoparticle-based visual detection can be done to increase the sensitivity of detection further. When aggregating agent, NaCl is added, the silver nanoparticles aggregate and produce grey colored solution in the absence of DNA. If DNA is present in the solution, the negatively-charged DNA masks the aggregation of silver nanoparticles from chloride salts, due to their strong association with positively-charged silver nanoparticles, and hence the solution remains yellow.

We first assessed the sensitivity or limit of detection using agarose gel electrophoresis employing ethidium bromide dye. Agarose gel electrophoresis of PCR products obtained using *esxA* and *mce3A* primers exhibit a sensitivity of detection of 4X10^4^ *M. tb* bacilli (~200pg of genomic DNA) **(Figure 4 A and B)**. In comparison to *M. tb*, *mmpS6* PCR is able to detect up to 4X10^3^ *M. bovis* bacilli (~20pg of *M. bovis* genomic DNA) **(Figure 4 C)**.

**Figure 4.**
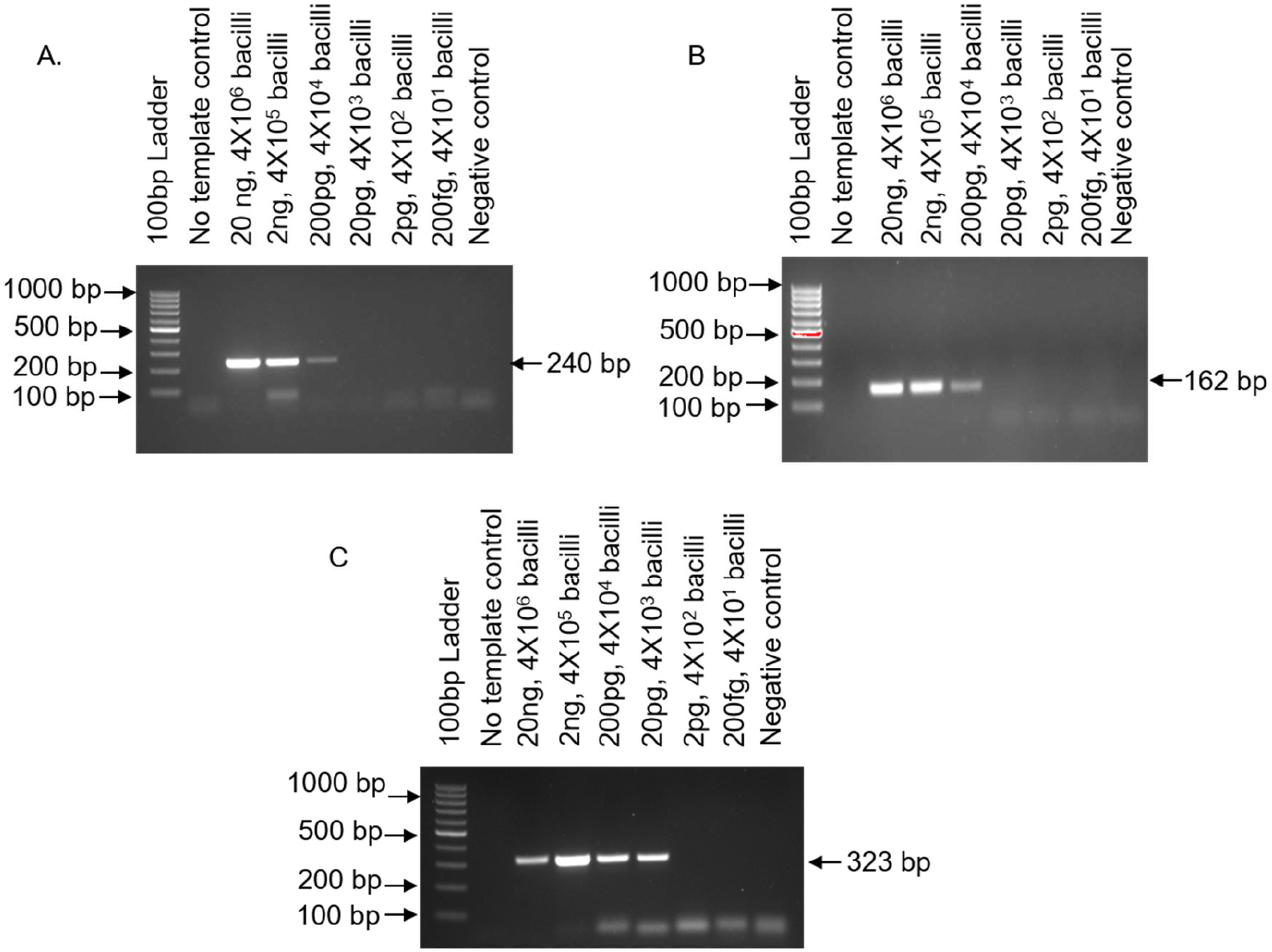
Sensitivity of *M. tuberculosis* and *M. bovis* PCR followed by agarose gel electrophoresis-based detection. The figure depicts agarose gel electrophoresis-based detection of PCR products stained with ethidium bromide dye. The labels on top of the gel indicate amount of genomic DNA used per PCR reaction and the corresponding number of bacilli. A. *esxA* gene-based detection of *M. tuberculosis.* B. *mce3A* gene-based detection of *M. tuberculosis.* C. *mmpS6* gene-based detection of *M. bovis. esxA* and *mce3A* PCR followed by agarose gel electrophoresis shows a detection sensitivity of ~200 pg of *M. tb* genomic DNA, corresponding to 4X10^4^ bacilli. *mmpS6* PCR showed a sensitivity of ~20 pg of *M. bovis* genomic DNA, corresponding to 4X10^3^ bacilli. No template control (NTC) and a mix of genomic DNAs except that of the test organism(s) served as negative control(s). For details see methods. (ng-nanogram; pg-picogram; fg-femtogram)

Paramagnetic beads are currently replacing the traditional silica-based spin columns for DNA purification post PCR-amplification due to ease, cost, scalability and turn-around time. In this study, we repurposed the paramagnetic beads for their capability to flocculate in the presence of DNA molecules obtained on PCR amplification. DNA molecules get typically adsorbed onto the surface of the paramagnetic beads, and addition of flocculation buffer containing sodium acetate (NaAc) and tween 20 at an acidic pH results in formation of DNA bridges connecting the paramagnetic beads, called as flocculation. However, in the absence of DNA or lesser amount of the amplified product, bridges are not formed indicated by brown dispersion with no floc formation ^25, 39^. Based on this assay, it was observed that, *esxA, mce3A* and *mmpS6* PCRs showed a detection sensitivity of ~40 bacilli of *M. tb.* and *M. bovis* **(Figure 5 A)**. Hence, this assay is far more sensitive than the traditional agarose gel based-DNA detection described above.

**Figure 5.**
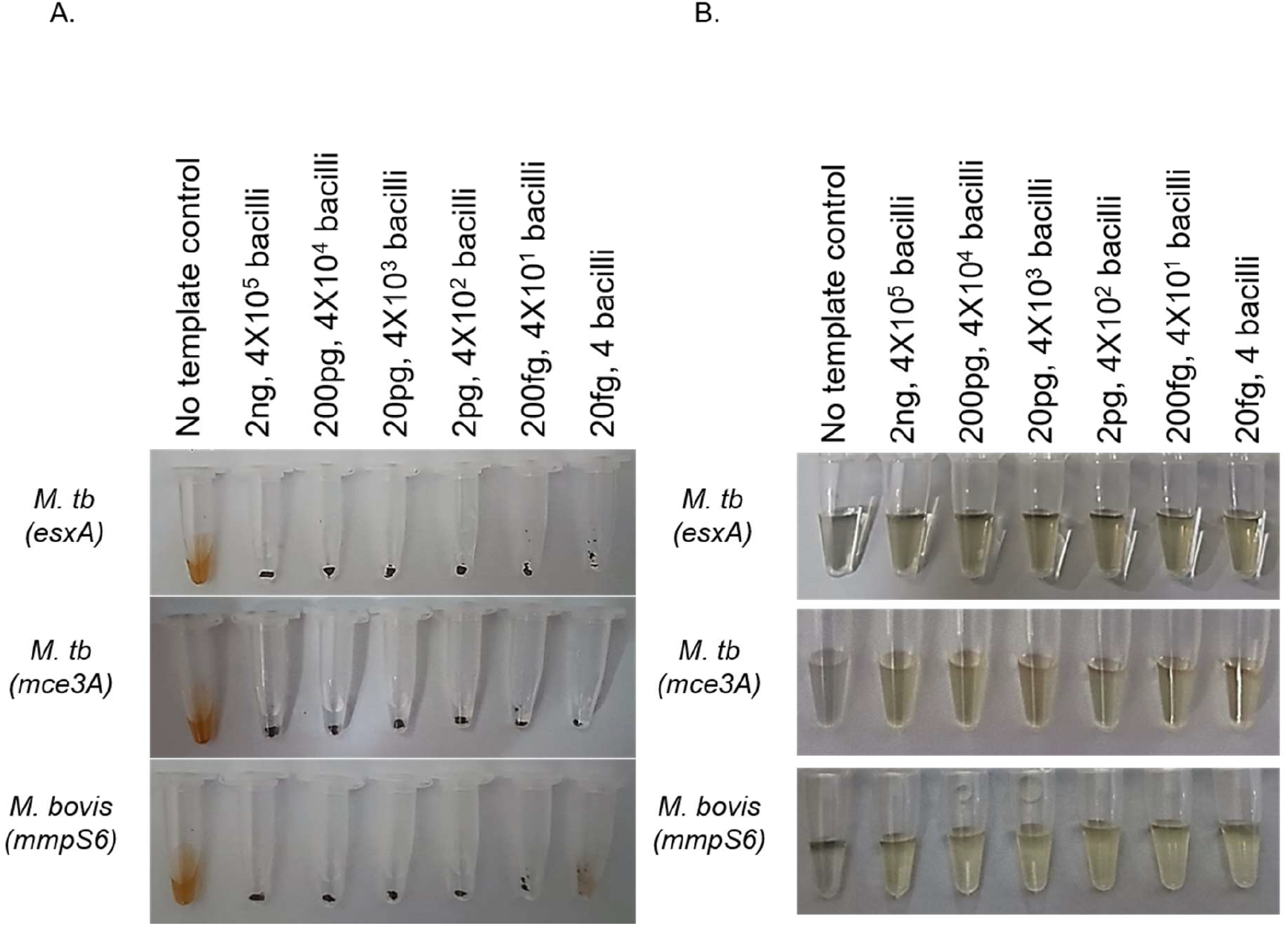
Sensitivity of paramagnetic bead flocculation and silver nanoparticle-based visual detection of *M. tuberculosis* and *M. bovis*. A. Paramagnetic bead-based bridging flocculation assay shows a uniform detection sensitivity of 200 fg of *M. tuberculosis* genomic DNA (~40 bacilli) utilizing *esxA* and *mce3A* gene specific primers and 200 fg of *M. bovis* genomic DNA (~40 bacilli), using *mmpS6* gene specific primers. Presence of PCR amplified DNA results in a visual floc (bead aggregate), however, absence or very low amount of amplified DNA result in dispersed state of magnetic beads (brown solution). B. Silver nanoparticle aggregation-based detection showed a detection sensitivity of ~4 bacilli in case of all three genes. Presence of amplified DNA causes silver nanoparticles to remain in dispersed state indicated by yellow color solution, however, absence or very low amount of amplified DNA results in aggregation of silver nanoparticles in presence of aggregating agent (NaCl), indicated by grey color solution. (ng-nanogram; pg-picogram; fg-femtogram)

### Citrate-stabilized AgNPs for visual detection of *M. tb* and *M. bovis*

Citrate-stabilized AgNPs have seldom been used for DNA detection. Here, we present for the first time a novel citrate-stabilized AgNP aggregation-based, label-free, visual detection of amplified DNA and compare the sensitivity of this technique with respect to traditional and widely researched agarose gel-based electrophoresis and bridging flocculation assay. The visual detection using these citrate-stabilized AgNPs is based on colour conversion from yellow to grey in the absence of DNA upon addition of high ionic strength aggregating agent such as NaCl. This happens due to chloride induced AgNP aggregation ^40^. However, when amplified & purified double stranded DNA (dsDNA) amplicons are present in the silver dispersion, the ionic association of negatively charged dsDNA with positively charged AgNPs prevent them from aggregating even on addition of NaCl ^41, 42^. This property of the AgNP aggregation has been utilized here to develop a highly sensitive visual detection method post DNA amplification. Our study reveals a detection sensitivity of 4 bacilli (~20 femtograms, fg) of mycobacterial genomic DNA) for both *M. tb*. and *M. bovis* using AgNPs **(Figure 5 B)**, which is far superior to agarose gel electrophoresis (4,000 - 40,000 bacilli) **(Figure 4 A-C)** and bridging flocculation assay (40 bacilli) **(Figure 5 A)**.

### Spectral characteristics of citrate-stabilized AgNPs in the presence of tris-EDTA buffer, dsDNA, and NaCl

In order to understand the underlying changes in aggregation profile of the AgNPs, we carried out a spectral analysis (300 – 700 nm) of silver nanoparticles in the presence of dsDNA of various amounts, tris-EDTA buffer and aggregating agent, NaCl. Aggregation of silver dispersion is characterised by reduction in the absorbance at ~396 nm, simultaneously, we observe an increase in the absorbance beyond 500 nm indicating the formation of large size AgNP aggregates. We observe that in comparison to, tris-EDTA buffer, NaCl acts as a relatively stronger aggregating agent, as evident from a higher reduction in A_396 nm_ on addition of NaCl **(Figure 6 A and B)**. Further, we observe a marked increase in A_520 nm_ on addition of NaCl **(Figure 6 A and C)**. However, in the presence of various amounts of dsDNA, no evident change is observed in the A_396 nm_ values, indicating arrival of the isosbestic point **(Figure 6 D and E)** ^41^. On addition of NaCl, to silver dispersion containing only tris-EDTA buffer, a markedly reduced A_396 nm_ is observed, however, in presence of various amounts of dsDNA, we observe a much higher A_396 nm_ commensurate to the amount of dsDNA **(Figure 6 F and G)**. These observations suggest that higher the amount of dsDNA, greater is the resistance towards aggregation causing the AgNP to remain in yellow colour. Interestingly, compared to visual changes in AgNP colour (grey vs yellow) which is more qualitative in nature with sensitivity up to 4 bacilli, spectral changes are far more sensitive in detecting the changes in yellow coloration and hence allows quantitative estimation of dsDNA.

**Figure 6.**
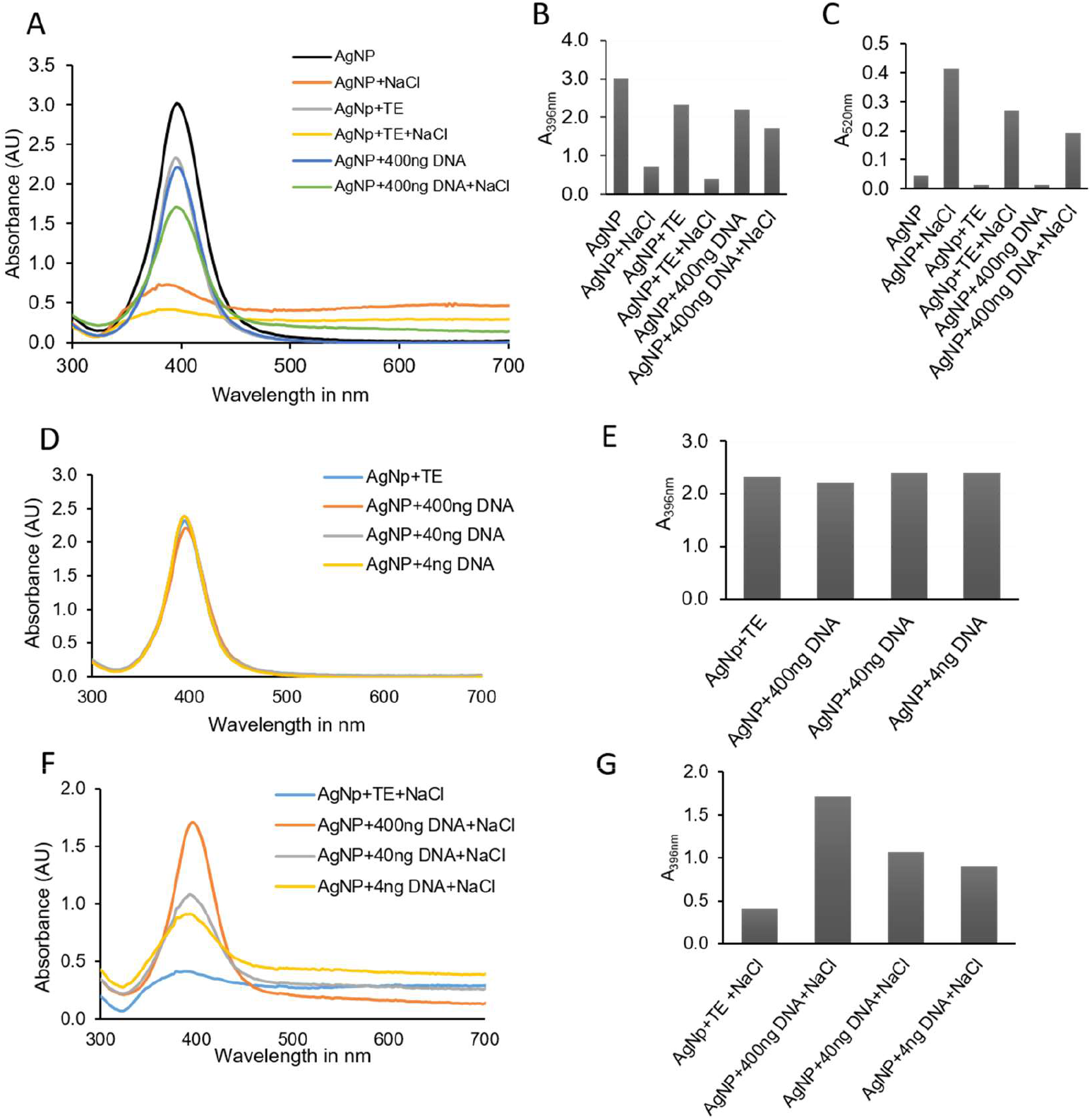
Spectral characteristics of citrate-stabilized silver nanoparticles (20nm) in the presence of tris-EDTA buffer, dsDNA, and NaCl. (A to C) Figure depicts increasing amount of aggregation of AgNPs characterized by reduction in A_396nm_ and an increase in A_520nm_. Tris-EDTA leads to reduction in A_396nm_ (indicating slight AgNP aggregation), whereas NaCl addition leads to a marked reduction in A_396nm_, signifying complete aggregation of AgNPs. Tris-EDTA buffer acts as a mild aggregating agent whereas, NaCl acts as a strong aggregating agent. Addition of negatively charged dsDNA, forms stronger ionic bonds with positively charged AgNPs, masking interaction with NaCl thereby preventing the aggregation of AgNPs on addition of NaCl. (D and E) With increasing amount of dsDNA, the spectral characteristics of AgNPs do not change leading to isosbestic point. (F and G) With decreasing amount of DNA, A_396nm_ reduces signifying increased AgNP aggregation on addition of NaCl. (nm – nanometre; ng – nanogram)

## Conclusion

In this study, we present a novel highly sensitive citrate-stabilized AgNP aggregation-based, label-free, visual method for differential detection of *M. tb* and *M. bovis* DNA. To the best of our knowledge, our study is the first to report application of citrate-stabilized silver nanoparticles for molecular detection of a pathogenic agent. Our study provides conclusive evidence for a far superior sensitivity of AgNP based detection (4 bacilli) compared to the existing agarose gel electrophoresis (4,000 – 40,000 bacilli). We utilize this highly sensitive method for differential detection of *M. tb* and *M. bovis* using specific primers designed to target unique regions of each of these mycobacterial species. We also compared sensitivity of AgNP-based detection with paramagnetic bead-based bridging flocculation assay for qualitative visual detection of *M. tb* and *M. bovis*. Our study is the first report for *M. bovis* detection using paramagnetic bead-based bridging flocculation assay and this method is found to be highly sensitive with detection limit of 40 bacilli. The AgNP and bridging flocculation methods for molecular detection reported by us in this study are far superior compared to the traditional agarose gel electrophoresis in terms of cost, time, sensitivity of detection, need of equipment for visualization and environmental impact **(Table 2)**.

**Table 2.** Comparative analysis of different detection techniques used in this study for the molecular identification of *M. tuberculosis* and *M. bovis*.

|  | **Agarose gel detection** | **Magnetic Bead Detection** | **Silver nanoparticle detection** |
| --- | --- | --- | --- |
| **Cost of detection (Excluding cost of PCR reagents and machine)** | Very High<br>(~\$ 6.5-10 including cost of chemicals, dyes, DNA markers, electrophoresis and gel casting accessories and electricity. Per sample cost excludes the required one-time investment for gel electrophoresis unit, power pack, gel documentation equipment and printer) | Very Low<br>(~\$ 1.5/sample including cost of paramagnetic beads, chemicals and plasticware. This test requires a one-time investment of a very cost-effective magnet stand. No cost for electricity or high-end equipment) | Low<br>(~\$ 2.0/sample inclusive of cost of paramagnetic beads and silver nanoparticle dispersion, chemicals and plasticware. This test requires a one-time investment of a very cost-effective magnet stand. No cost for electricity or high-end equipment) |
| **Need of equipment for visualization** | UV Gel documentation equipment for visualisation. | Naked eye | Naked eye |
| **Time to detect (Post amplification)** | 2- 2.5 hours<br>(No DNA purification step) | 15-20min<br>(Simultaneous DNA purification and visual detection) | 20-30min (DNA purification followed by silver nanoparticle based visual detection) |
| **Sensitivity (number of detectable bacilli)** | Less sensitive<br>(4000 – 40,000 bacilli) | High sensitivity<br>(40 bacilli) | Very high sensitivity<br>(4 bacilli) |
| **Environment impact** | Environment and user non-friendly, unsafe, utilizes carcinogenic agents such as Ethidium bromide and UV. | Non-carcinogenic, environment and user friendly | Non-carcinogenic, environment and user friendly |
| **Ease of performance** | Difficult to perform, require highly skilled technical personnel and controlled environment needing continuous supply of electricity | Easy to perform and analyze by less skilled personnel without the need of controlled environment and electricity | Easy to perform and analyze by less skilled personnel without the need of controlled environment and electricity |
| **Individual testing** | Single sample testing is not cost-effective | Single sample can be tested individually | Single sample can be tested individually |

Citrate-stabilized AgNPs have several merits like greater stability than non-stabilized AgNPs against photo-oxidation, ensuring a reliable and consistent signal over time, which enhances the robustness of the detection method ^43^. AgNPs are cost-effective compared to the gold nanoparticles, making them a practical choice for DNA detection assays in various settings. The relevance of citrate-stabilized silver nanoparticles for DNA detection lies in their unique properties that make them suitable for developing sensitive and specific nucleic acid detection assays ^44, 45^. The colorimetric response of citrate-stabilized silver nanoparticles, resulting from changes in plasmon resonance upon DNA hybridization or target DNA binding, further provides a visual and straightforward means of detection without the need for complex instruments. Overall, the silver nanoparticle-based DNA detection offers a versatile method, making it a promising alternative for diagnostic and research applications. Methods reported here are also easy to perform by less skilled personnel and are compatible with individual testing, hence can be performed in POC settings with PCR machines. The label-free silver nanoparticle aggregation approach, if coupled with isothermal amplification methods like Loop-mediated Isothermal Amplification (LAMP) ^46^ and Recombinase Polymerase Amplification (RPA) ^47–52^, could revolutionize POC diagnostics, shifting tuberculosis management. Combining these techniques could also markedly reduce detection costs, time, and procedural complexities inherent in TB diagnosis. The visual detection could possibly be utilized for the molecular detection of drug resistant forms of *M. tb* or *M. bovis* ^51–53^. However, it is crucial to acknowledge that visual detection methods may not provide quantitative results, which can be important for monitoring disease progression and treatment response.

## Materials and methods

### Ethics Statement

The study was performed as per the guidelines and approval of the institutional biosafety committee (BITS/IBSC/2018-2/04).

#### Reagents and Resources

All the reagents used for the study were procured from Himedia Laboratories Pvt. Ltd., unless stated otherwise. The oligonucleotide primers used for the study as described in Table 1, were procured from Sigma Aldrich. Paramagnetic beads were procured from MagGenome. Silver nanoparticles (citrate buffer stabilised, 20 nm size, 20 mg/ml) were procured from Sigma Aldrich. *Taq* polymerase and their supporting reagents were procured from Takara Clontech. *Neisseria gonorrhoeae (*ATCC 19424) was purchased from ATCC (Himedia Laboratories Pvt. Ltd.). WHO BCG Danish 1331 vaccine sub-strain was purchased from National institute for Biological Standards and Control (NIBSC), UK.

#### PCR-based molecular detection

For *M. tb* detection, we curated the oligonucleotide primers from published literature ^24, 33^ and for *M. bovis* we designed the primers in this study using the PrimerQuest tool (Integrated DNA Technologies). Primers were designed to keep the amplicon size within the range of 150 bp – 500 bp (Table 1). The specificity of the primers was verified *in-silico* using NCBI-BLAST. PCR was performed in Veriti Thermal cycler, Applied Biosystems, using *Taq* Polymerase from Takara Clontech, as per the manufacturer’s instructions with slight modifications. 50 μl PCR reactions were set up with 1 μM primer mix in each reaction mixture. Thermal cycling was performed with an initial denaturation of 95°C for 10 minutes and 40 cycles of each denaturation (45 seconds at 95°C), annealing (45 seconds at the optimized annealing temperature), extension (1 minute at 72°C), and final extension at 72°C for 10 minutes. The optimum annealing temperatures for the respective primers were deciphered using temperature gradient PCR. In order to validate specificity of the primers towards target organism(s)-*M. tb* and *M. bovis*, a mix of genomic DNA obtained from a variety of other organisms such as, *M. bovis* BCG, *Neisseria gonorrhoeae*, *Chlamydia trachomatis*, *Ureaplasma urealyticum,* and human were used as a negative control sample. Agarose gel-based DNA electrophoresis was used for verification of PCR products. In order to determine the sensitivity or limit of detection, varying amounts of the genomic DNA template obtained from respective organisms were used for performing PCR and verifying the same by agarose gel electrophoresis employing ethidium bromide dye for visualization. Total procedure of gel preparation, electrophoresis and detection takes ~2-2.5 hours.

#### Visual detection using bridging flocculation assay

Bridging flocculation assay was performed using paramagnetic beads from MagGenome. 20 μl of the PCR product was volume adjusted to 50 μl using sterile distilled water. Equal volume of the paramagnetic bead solution was added to the PCR product and incubated at room temperature for 10-15 minutes. The beads were then magnetically separated and washed with 80% ethanol (wash buffer) on the magnetic stand (MagGenome). 100 μl of flocculation buffer (200 mM sodium acetate, pH 4.4, 1% v/v Tween 20) was added to the washed beads. The tubes were then incubated in the flocculation buffer for 1-2 minutes on the magnetic stand. The tubes were then removed from the stand and gently tapped at bottom until the beads in the no-template-control (NTC) were dispersed into a brown solution. Tubes with amplified product were similarly tapped, however, these tubes showed a visual floc, resistant to dispersion on gentle tapping. Total procedure of purification and visual detection takes 15-20 minutes. Sensitivity of detection was determined by carrying out flocculation assay with amplicons obtained from PCR conducted with varying amounts of genomic DNA obtained from target organisms *M. tb* and *M. bovis*.

#### Visual detection using citrate-stabilized silver nanoparticles (AgNPs)

PCR products (20 μl) were cleaned-up using MagGenome PCR clean up kit as per the manufacturer’s instructions. The elution was done in 20 μl of elution buffer (10 mM Tris, 1mM EDTA, pH 8.0). 50 μl of the citrate-stabilized silver nanoparticle dispersion was added to 20 μl eluent and incubated at room temperature for a minute. 5 μl of aggregating reagent (5M NaCl) was added to the tubes. AgNP aggregation is immediately indicated by grey colour dispersion in tubes without amplicons (NTC or negative controls containing DNA of non-target organisms), whereas in tubes with amplified DNA, AgNPs do not aggregate on NaCl addition, hence the dispersion remains yellow. Total procedure of purification and visual detection takes 20-25 min. Sensitivity of detection was determined by carrying out citrate-stabilised AgNP aggregation based-assay with amplicons obtained from PCR conducted with varying amounts of genomic DNA obtained from target organisms *M. tb* and *M. bovis*.

#### Spectral analysis of AgNP in presence of dsDNA and aggregating agent NaCl

The spectrum was obtained in an Eppendorf Biospectrometer in a wavelength range of 300 nm to 700 nm. Varying amounts of purified DNA amplicons in tris-EDTA buffer (400 ng, 40 ng, and 4 ng) in 20 µl were mixed with 50 µl of silver dispersion and the respective spectra were obtained. Silver dispersion alone and in presence of elution buffer (10 mM Tris, 1 mM EDTA, pH 8.0) were used as the controls in this assay. Spectral analysis was also performed in the negative controls as well as test samples after addition of 5 µl of aggregating agent (5M NaCl).

#### Data Analysis

The data obtained from spectral analysis were plotted using MS Excel and spectra and histograms were deduced.

## Ancillary Information

### Corresponding Author Information

Ruchi Jain Dey -

### Author Contribution

NP and RJD have conceived and designed the project, analysed the results, designed figures and written and edited the manuscript. RJD and NP received the funding.

## Acknowledgements

RJD is thankful to Birla Institute of Science and Technology (BITS) Pilani, Hyderabad campus, India for their funding support through intramural funding under Research Initiation Grant (RIG) and Centre for Human Disease Research (CHDR). RJD is also thankful to the Department of Biotechnology (DBT) for supporting research through Ramalingaswami Re-entry fellowship (BT/RLF/Re-entry/18/2016). RJD is thankful to the overall infrastructure support by Department of Biological Sciences, BITS Pilani Hyderabad. NP is thankful to Indian Council of Medical Research (ICMR), Govt. of India for providing Senior Research Fellowship (RBMH/FW/2019/13). Dr. Raghunand R Tirumalai, Centre for Cellular and Molecular Biology (CCMB), Hyderabad, India is acknowledged for providing genomic DNA for *M. tuberculosis H37Rv*. Dr. Bappaditya Dey, National Institute of Animal Biotechnology (NIAB), Hyderabad, India is acknowledged for providing genomic DNA for *M. bovis.* We are thankful to Dr. Benu Dhawan, Department of Microbiology, All India Institute of Medical Sciences (AIIMS, New Delhi, India) for providing genomic DNA from a clinical isolate of *Ureaplasma urealyticum*. Dr. Karthika Rajeeve, Rajiv Gandhi Centre for Biotechnology, Trivandrum, India is highly acknowledged for providing genomic DNA of *Chlamydia trachomatis* (L2 Serovar).

## Data availability statement

All data that supports the findings of this study are available from corresponding author upon reasonable request.

## Abbreviations

*M. tb*: *Mycobacterium tuberculosis*
*M. bovis*: *Mycobacterium bovis*
TB: Tuberculosis
PTB: Pulmonary tuberculosis
EPTB: Extra-pulmonary tuberculosis
DOTS: Directly Observed Treatment, Short Course
MTBC: *Mycobacterium Tuberculosis* Complex
bTB: Bovine TB
POC: Point-of-care
AFB: Acid-Fast Bacilli
ml: Millilitre
PCR: Polymerase Chain Reaction
DNA: Deoxyribonucleic acid
qPCR: Quantitative PCR
PZA: Pyrazinamide
IS6110: Insertion Sequence 6110
ITS: Internal Transcribed Spacer
TbD1: Tb Deletion Region 1
NTC: no-template-control
NaAc: Sodium Acetate
AgNP: Silver Nanoparticles
dsDNA: Double stranded DNA
Tris: Tris(hydroxymethyl)aminomethane
EDTA: Ethylene diamine tetra-acetic acid
NaCl: Sodium Chloride
LAMP: Loop-mediated Isothermal Amplification
RPA: Recombinase Polymerase Amplification
BCG: Bacille Calmette-Geurin
WHO: World Health Organisation.

